# Receptor Activity-Modifying Protein 3 enhances GLP-1-mediated Insulin Secretion

**DOI:** 10.1101/2025.01.24.634724

**Authors:** Abigail Pearce, Poonam Kumari, Claudia M Sisk, Matthew Harris, Ho Yan Yeung, Sabrina Winfield, Kathleen M Caron, Graham Ladds

**Affiliations:** Department of Pharmacology, University of Cambridge, Tennis Court Road, Cambridge, CB2 1PD, UK; Department of Cell Biology and Physiology, School of Medicine, University of North Carolina at Chapel Hill, Chapel Hill, NC 27599

**Keywords:** G Protein-coupled Receptor, Allosteric Regulation, Diabetes, Second Messenger, insulin secretion, Calcium intracellular release

## Abstract

The targeting of the Glucagon-like peptide-1 receptor (GLP-1R) for diabetes and obesity is not a novel strategy, with recent therapeutics showing efficacy in weight loss and glycaemic control. However, they are also associated with side effects, including gastrointestinal disruptions and pancreatitis. Developing agonists with different signalling profiles, or that exert some tissue selectivity can circumvent these on-target, unwanted effects. Receptor activity-modifying proteins (RAMPs) offer the potential to do both, through modulation of agonist binding and signalling, as well as surface expression. The GLP-1R was found to interact with RAMP3, with the heterodimer able to bind agonist at the cell surface. RAMP3 expression biased the receptor towards Ca^2+^ mobilisation, away from the canonical cAMP-driven signalling. When examining G protein coupling, the interaction with RAMP3 reduced activation of the cognate Gα_s_ but increased secondary couplings to Gα_q_ and Gα_i_. These increased couplings led to an elevation in glucose-stimulated insulin secretion when cells overexpressing RAMP3 were stimulated with GLP-1. A reciprocal effect was observed when looking at reduced expression of endogenous RAMP3, with a loss of sensitivity to GLP-1 in both glucose and insulin tolerance tests in a *Ramp3* knockout mouse. The effects of this interaction can then inform selection of models and peptide design when targeting this receptor for therapeutic intervention.

**Significance Statement:** G Protein-Coupled Receptors, such as the Glucagon-Like Peptide-1 (GLP-1) Receptor are common drug targets, although not without side effects due to on-target, unwanted signalling outputs. This study identifies an interacting protein which alters its signalling to promote insulin secretion. This may improve the therapeutic potential of GLP-1 mimetics, enhancing effectual signalling over that associated with unwanted effects by directly targeting this receptor complex.

## Introduction

Obesity and type 2 diabetes mellitus (T2DM) are increasingly prevalent diseases, with the incidence expected to rise to 51% of the global population by 2035 (https://data.worldobesity.org/publications/?cat=19). Finding treatments for these diseases has been an ongoing mission, with efforts more recently focused on simultaneous targeting of both conditions, with the aim to improve T2DM by resolving the underlying obesity.

A recent therapeutic advance has been the approval of peptides targeting the Glucagon-Like Peptide-1 (GLP-1) receptor for obesity and weight loss. The incretin effect, whereby oral glucose elevates plasma insulin levels to a greater degree than IV administration, was first observed in 1964 (1). This is mediated by incretin hormones, the best known of which are GLP-1 and the glucose-dependent insulinotropic polypeptide (GIP). As such, mimetics of the endogenous GLP-1 hormone, the first of which was exenatide (exendin-4), were approved for T2DM in 2005 (2). However, newer mimetics semaglutide (trade names of Ozempic and Wegovy) or tirzepatide (sold under Mounjaro or Zepbound), the latter of which targets both the GLP-1R and the GIP Receptor, have garnered much attention through their dual effects on lowering plasma glucose and aiding weight loss. Semaglutide has since been reported to protect against neurodegeneration and neuroinflammation (3), as well as cardiovascular disease (4). This culminated in semaglutide being labelled as the 2023 breakthrough of the year (5). However, with their increased use, attention is being drawn to the unwanted side effects of GLP-1 mimetics. Common side effects are nausea and vomiting, as well as other gastrointestinal discomforts. However, more serious side effects are also known, such as pancreatitis, kidney failure, and gallbladder problems (6). There is therefore the unmet need to develop mimetics with improved therapeutic potential that exhibit fewer side effects.

One method of reducing on-target unwanted effects is to selectively activate signalling pathways associated with the positive outcome, relying on the concept of signalling bias. The GLP-1R is a class B1 G protein-coupled receptor (GPCR), which predominantly couples to Gαs, leading to an increase in cyclic-adenosine monophosphate (cAMP) production. However, like other members of class B1, the GLP-1R exhibits pleiotropy, coupling to other Gα families and recruiting β-arrestins (7, 8), leading to a multifaceted signalling output. For other pleiotropic GPCRs, the activation of these different pathways can be influenced by the agonist, giving rise to signalling bias. This has been studied most heavily for the μ-opioid receptor, where it is thought agonist tolerance is mediated by β-arrestins, leading G protein-biased agonists to be preferred (9).

Signalling bias can also be influenced by the tissue, due to differential expression of signalling proteins, as well as allosteric modulators. A common protein allosteric modulator of Class B1 GPCRs are the receptor activity-modifying proteins (RAMPs). First discovered for the calcitonin-like receptor, with which they form obligate heterodimers, RAMPs influence the expression, affinity, and signalling bias of the receptor. For the calcitonin family, this leads to receptors considered functionally distinct (10, 11), although smaller effects are observed on other receptors e.g. the vasoactive intestinal peptide receptor 1 (VPAC_1_) or glucagon receptor (GCGR) when co-expressed with RAMP2 (12). Importantly, the effects of RAMPs are agonist- and pathway-dependent, with RAMP2 increasing Gα_s_ coupling at GCGR in response to glucagon and oxyntomodulin (an endogenous dual agonist of both GCGR and GLP-1R) but decreasing the response to GLP-1 and liraglutide (a synthetic, lipidated GLP-1 mimetic, approved in the treatment of T2DM) (13).

RAMPs have been implicated in diabetes and obesity, although their role appears multifaceted. Of the GPCRs which interact with RAMPs, many are involved in the regulation of glucose levels and body weight. The pancreatic hormone amylin, binds a class B1 GPCR, with the three amylin receptors heterodimers between the calcitonin receptor (CTR) and RAMP1-3. RAMPs are expressed in many of the tissues of the endocrine system, such as the thyroid and hypothalamus, and global RAMP knock out in rodents causes dysregulation of body weight and glycaemic control (14, 15) (the role of RAMPs in diabetes and obesity is further reviewed in Malcharek et al., 2024 (16)). Importantly, RAMPs are co-expressed with the incretin hormone receptors across regions of the brain, and the pancreas (Human Protein Atlas as of January 2025). The interaction between GIPR and RAMPs, and the consequences for glucose tolerance have been studied previously (17), but the involvement of GLP-1R is less known. GLP-1R has been previously shown to interact with all three RAMPs (18, 19), with depressive effects on cAMP accumulation observed. However, specific interactions at the plasma membrane have not been studied, nor have any effects of RAMP expression on physiological roles of the receptor, such as glucose-stimulated insulin secretion (GSIS).

Herein, we measured a specific membrane interaction between GLP-1R and RAMP3, but not RAMP1 or 2. The two proteins were observed to form a complex able to bind agonists, but no effect was observed on affinity of the receptor. RAMP3 expression decreased cAMP accumulation and increased mobilisation of Ca^2+^ from intracellular stores (Ca^2+^)_i_. This was determined to be primarily due to an increase in Gα_i/o_coupling. In contrast, no effect was observed on internalisation, with only a decrease in maximal β-arrestin recruitment. This elevation in (Ca^2+^)_i_ was shown to lead to enhanced GSIS. The reciprocal effect was observed in murine cell and animal models, where a knockdown or knockout of *Ramp3* decreased insulin secretion in response to GLP-1. There is therefore the potential for improved GLP-1 mimetics, which show bias towards the GLP-1R-RAMP3 complex, with subsequent effects on tissue selectivity and therapeutic profile.

## Results

### GLP-1R specifically interacts with RAMP3

Although GLP-1R expression has been unable to promote RAMP membrane trafficking, interactions have been observed at the molecular level, e.g. using BRET (19). Therefore, to determine if these observed interactions are present at the plasma membrane, a cell surface BRET interaction assay was used as previously described (17). Nluc-GPCR and SNAP-RAMP were transiently transfected into Cos7 cells, which do not endogenously express RAMPs, CTR or CLR (20), to reduce any interference from unlabelled proteins. SNAP-RAMPs were irreversibly conjugated to SNAP-Surface Alexa Fluor 488 to specifically measure cell surface interactions; the conjugate was cell impermeable so only labelled SNAP-RAMPs expressed at the cell surface. All three RAMPs displayed an interaction with CLR, as observed by the saturating increase in BRET ratio. GLP-1R only displayed a saturating increase in BRET with SNAP-RAMP3 (Figure 1A). Overexpression of HA-CLR was not shown to promote an interaction between Nluc-GLP-1R and SNAP-RAMP1 or 2, indicating that expression of RAMP at the membrane is not sufficient to induce an interaction, even when increasing the amount of SNAP-RAMP ten-fold over Nluc-GLP-1R.

**Figure 1.**
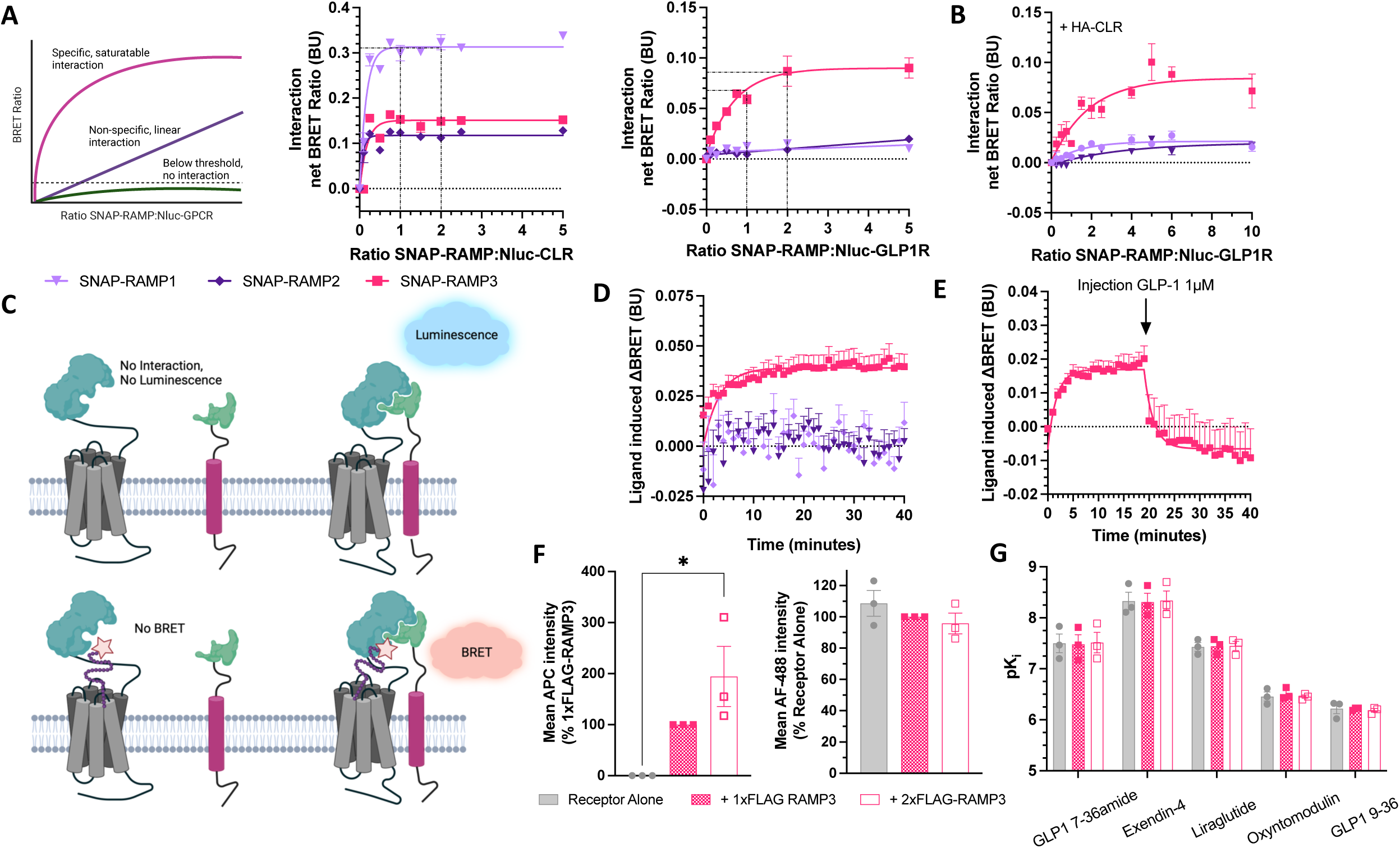
GLP-1R and RAMP3 interact and can bind agonists as a complex. (A) Cell-surface NanoBRET assay measuring interactions between Nluc-CLR (*left*) and Nluc-GLP-1R (*right*) with SNAP-RAMPs labelled with SNAP-Surface Alexa Fluor 488 in Cos7 cells, n=4. (B) Nluc-GLP-1R interaction with labelled SNAP-RAMPs, in the presence of a fixed concentration of HA-CLR, n=8. (C) Schematic of a NanoBiT assay for measuring GLP-1R-RAMP interactions and fluorescent ligand binding to the complex. (D) Binding of Ex-4-Red to the LgBiT-GLP-1R SmBiT-RAMP3 complex in HEK293T cells (n>4), which can then be displaced by addition of 1μM unlabelled GLP-1 (E), n=3. (F) Expression of FLAG-RAMP3 (*left*) or Nluc-GLP-1R (*right*) in HEK293T cells transfected with Nluc-GLP-1R and increasing amounts of FLAG-RAMP, n=3. (G) Binding affinities (pK_i_) values for GLP-1, exendin-4, liraglutide, oxyntomodulin, and GLP-1 9-36 at Nluc-GLP-1, co-expressed with FLAG-RAMP3, n=3. Mean values of n experimental replicates are shown ± SEM, statistical differences determined through Kruskal-Wallis test for non-parametric data or One-Way ANOVA, *\** p < 0.05.

To validate the formation of a GLP-1R-RAMP3 complex, we measured binding of a fluorescent ligand to the protein heterodimer. We utilised NanoBiT, with LgBiT preceding the N-terminus of the receptor, and SmBiT upstream of the N-terminus of the RAMP (illustrated in Figure 1C). Binding of the Tag-Lite GLP-1 receptor red ligand, based on exendin-4 (Ex-4-Red) was observed through an increase in the BRET ratio, with a calculated K_d_ of 6.7nM (Figure 1D). This was not observed for SmBiT-RAMP1 or -RAMP2. The BRET ratio could be reduced by injection of 1μM unlabelled GLP-1, competing with the fluorescent agonist and further demonstrating binding had occurred to the heterodimer (Figure 1E).

### FLAG-RAMP3 has no effect on GLP-1R expression or agonist binding

When measuring their effects on GPCR signalling, RAMPs are commonly co-expressed at a 1:1 ratio, due to the stoichiometry of the interaction with CLR (21). However, the maximal interaction between GLP-1R and RAMP3 in the BRET assay was observed when RAMP3 was expressed at twice that of GLP-1R (by weight of expression plasmid, Figure 1A). We therefore measured the effect of increasing RAMP3 expression on GLP-1R, either co-expressing (1:1 ratio) or overexpressing (2:1 ratio) RAMP3. It was confirmed that the expression of FLAG-RAMP3 was increased at the 2:1 ratio compared to 1:1, but no difference was observed on GLP-1R expression (Figure 1F).

In addition to receptor expression, differences in signalling could be driven by changes in affinity. As RAMP modulation of GPCR signalling can be agonist dependent, effects were examined across a range of agonists. GLP-1 and oxyntomodulin are endogenous agonists, the latter with a reduced potency. GLP-1 9-36, a product of the degradation of GLP-1 which can show weak partial agonism, was also tested. Examples of incretin mimetics, exendin-4 and liraglutide, were also included. FLAG-RAMP3 had no influence on binding affinity for any peptide as measured through a NanoBRET competition binding assay (Figure 1G).

### Second messenger signalling is influenced by RAMP3 expression

As with most Class B1 GPCRs, the primary signalling mediator for the GLP-1R is Gα_s_, leading to an increase in cAMP (7, 22). When stimulated with GLP-1 for 8-minutes, a dose-dependent increase in cAMP was observed (pEC_50_ value of 11.4±0.3). This was reduced by RAMP3 expression, with a reduction in the potency at 8-minutes when RAMP3 was over expressed (to 10.4±0.2) (Figure 2A, Table 1). The pattern was preserved across exendin-4, liraglutide, and oxyntomodulin, with reductions in both potency and efficacy observed when RAMP3 was overexpressed (Figure 2B). GLP-1 9-36 did not elicit sufficient cAMP production over the concentration range tested to calculate potency and efficacy values.

**Figure 2.**
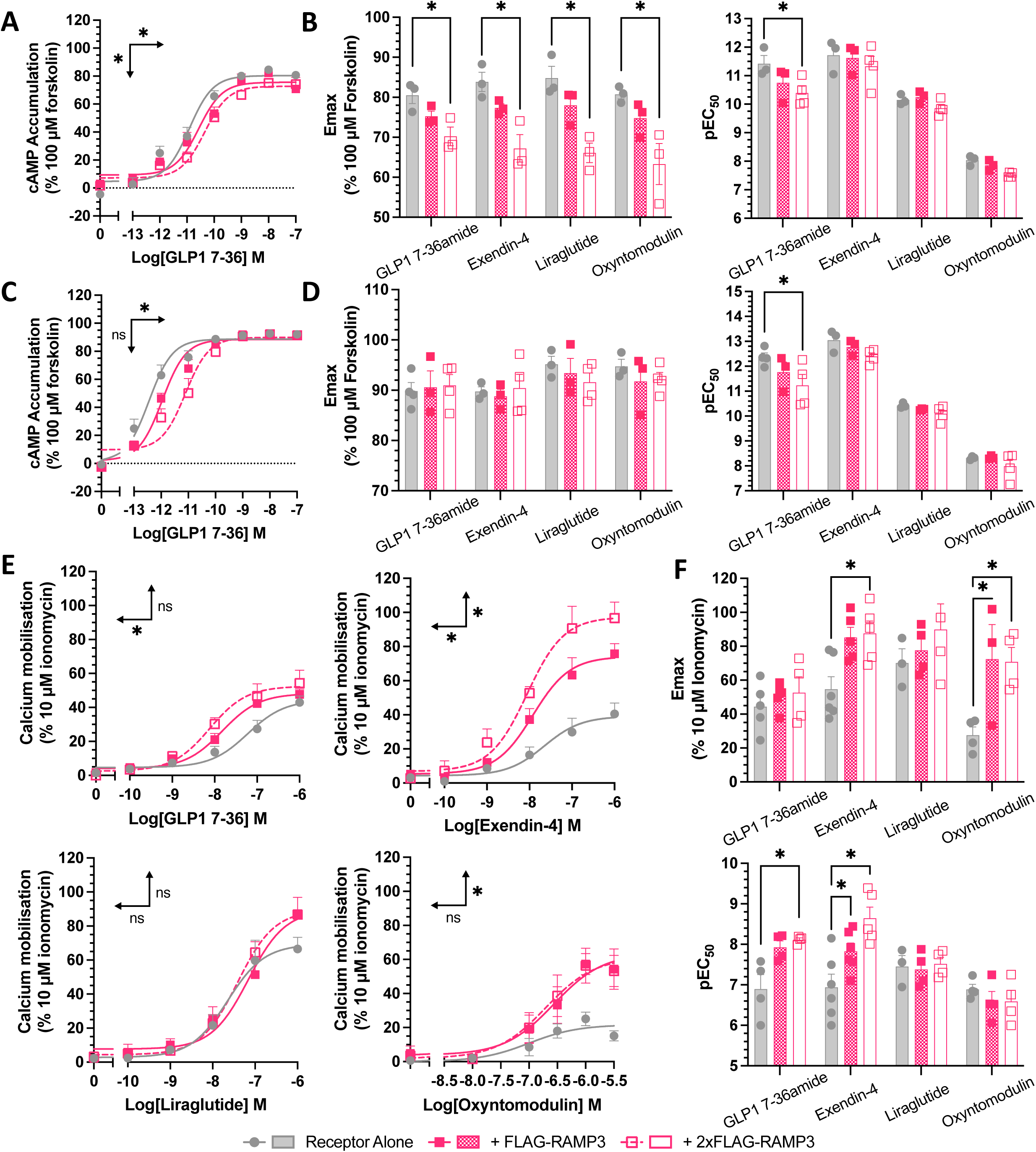
RAMP3 decreases cAMP accumulation but increases (Ca^2+^)_i_ mobilisation. cAMP accumulation mediated by Nluc-GLP-1R, expressed with increasing concentrations of FLAG-RAMP3, in response to 8-minute stimulation with GLP-1 (A), with E_max_ and pEC_50_ values plotted for GLP-1, exendin-4, liraglutide, or oxyntomodulin (B). (C-D) As before but for 30-minute stimulation. Responses are normalised to 100μM forskolin, n>3. (E) Mobilisation of (Ca^2+^)_i_ by Nluc-GLP-1R with FLAG-RAMP3, in response to stimulation with GLP-1, exendin-4, liraglutide, or oxyntomodulin, with E_max_ and pEC_50_ values plotted (F). Responses are normalised to 10μM ionomycin, n>3. Mean values of n experimental replicates are shown ± SEM, significance between RAMP conditions determined through One-Way ANOVA, * p < 0.05. Horizontal arrows indicate changes in potency (pEC_50_), and vertical designate changes in efficacy (E_max_).

When increasing the stimulation period to 30-minutes, there was again a reduction in the potency of the response to GLP-1, with over-expression significantly reducing the pEC_50_ value (Figure 2C, Table 1). Effects on other agonists were reduced, with significant differences no longer observed (Figure 2D).

GLP-1R is a pleiotropic receptor, able to couple to G proteins other than Gα_s_ (7, 8, 23). In many studies, the receptor has been found to couple to the Gα_q/11_ family of G proteins, with GLP-1R activity also associated with increased intracellular Ca^2+^ concentrations (Ca^2+^)_i_ due to mobilisation from intracellular stores. The effect of RAMP3 were therefore tested against (Ca^2+^)_i_ mobilisation (Figure 2E-F, Table 1). FLAG-RAMP3 expression enhanced the (Ca^2+^)_i_ mobilisation in response to GLP-1, increasing the potency of the response (Figure 2E). The dataset was again extended to other agonists to determine any agonist-dependent effects. Again, GLP-1 9-36 was unable to elicit any (Ca^2+^)_i_ mobilisation in the absence of RAMP3, meaning signalling parameters could not be calculated.

There was a significant trend for RAMP3 to increase maximal signalling in response to all agonists (*p<0.0001*) (Figure 2E-F, Table 1). Expression in a 1:1 ratio significantly increased the maximal responses to exendin-4 (from 43.6±5.30 to 71.61±4.64 percent of 10 μM ionomycin, *p<0.05*) and oxyntomodulin (from 21.7±3.5 to 63.6±10.6 percent of 10 μM ionomycin, *p<0.01*). Over-expression also significantly increased the maximal response of these two agonists to 97.6±6.8 and 52.3±8.5 respectively (*p<0.0001, p<0.05*), as well as increasing the potency of the response to GLP-1 (from a pEC_50_ of 7.2±0.2 to 8.1±0.2, *p<0.05*). There were no significant effects on the response to liraglutide, highlighting agonist dependency.

As changes in signalling can result from altered receptor desensitisation, we investigated the effects of RAMP3 expression on β-arrestin recruitment and internalisation. This is especially pertinent when concerning RAMP3, as it is known to have effects on receptor internalisation and trafficking for other class B1 GPCRs (24–26). Recruitment was reduced for all agonists except oxyntomodulin, although only when RAMP3 was expressed at a 2:1 ratio of receptor (Figure 3A). For GLP-1, exendin-4, and oxyntomodulin, there was no subsequent effect on internalisation, consistent with the observation that GLP-1R internalisation is β-arrestin-independent (27) (shown for GLP-1 in Figure 3B, Table 2). However, RAMP3 overexpression led to a significant decrease in the potency of internalisation in response to liraglutide (Figure 3C, Table 2).

**Figure 3.**
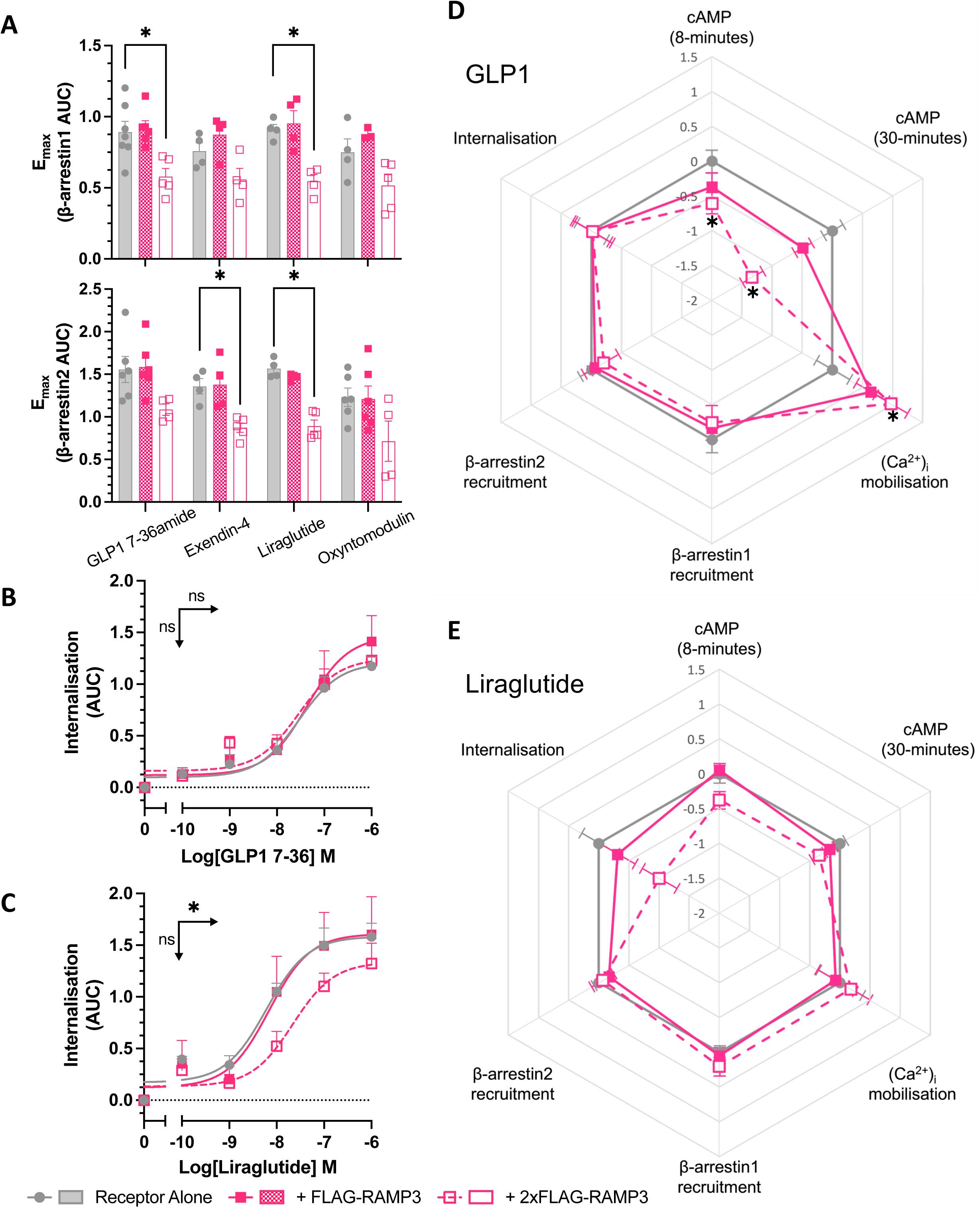
Effects of desensitisation are agonist dependent. (A) Maximal β-arrestin1 (upper) or β-arrestin2 (lower) recruitment to GLP-1R-Nluc, in response to GLP-1, exendin-4, liraglutide, or oxyntomodulin, n>4. Internalisation, as measured through a loss of BRET between GLP-1R-Nluc and plasma membrane marker RIT-Venus, in response to GLP-1 (B) or liraglutide (C), n>3. (D-E) LogRA values for second messenger signalling and receptor desensitisation for GLP-1R, measured in HEK293T cells, in response to GLP-1 or liraglutide (n>3). Responses are normalised to the response of GLP-1 at GLP-1R alone. Mean values of n experimental replicates are shown ± SEM, significance between RAMP conditions determined through One-Way ANOVA, * p < 0.05. Horizontal arrows indicate changes in potency (pEC_50_), and vertical designate changes in efficacy (E_max_).

In order to make comparisons across pathways and agonists, logarithmic values of relative activity (LogRA) were calculated as metrics of bias for second messenger signalling assays (cAMP and (Ca^2+^)_i_ mobilisation) and desensitisation (Tables 1-2). The effects of RAMPs overall varied when looking at different peptides. GLP-1 showed a decrease in cAMP accumulation, and an increase in (Ca^2+^)_i_ mobilisation, with little effect on desensitisation (Figure 3D). However, liraglutide displayed lesser effects on second messenger signalling, but a greater reduction in internalisation (Figure 3E).

### RAMP3 expression influences G protein coupling

To determine the underlying changes in G protein coupling associated with this biased second messenger production, the TRUPATH G protein biosensor platform was used (28). Three major families were examined, as they are closely linked with changes in cAMP accumulation and (Ca^2+^)_i_ mobilisation; Gα_s_, Gα_i/o_, and Gα_q/11_. There are reports that the TRUPATH system struggles to detect Gα_q_ coupling at classically Gα_s_ coupled receptors (28, 29). We therefore tested the wild type Gα_q_, and the R183Q mutant (Gα_q_(Q)), which displays increased constitutive and agonist-induced activity (30). in response to all four full agonists. No quantifiable response was observed using wild type Gα_q_, as has been observed previously (29).

Whilst no effect was observed for GLP-1, exendin-4, or liraglutide, there was a significant reduction in the maximal Gα_s_ dissociation elicited by oxyntomodulin when RAMP3 was expressed (reduced to 58.9% ±5.8 of receptor alone) (Figure 4A, Supporting Table S1). There was no effect on the Gα_q_(Q) mutant response to exendin-4, liraglutide, or oxyntomodulin, but RAMP3 did significantly increase the maximal dissociation elicited by GLP-1 (to 119 ±5.5% of receptor alone) (Figure 4B, Supporting Table S1).

**Figure 4.**
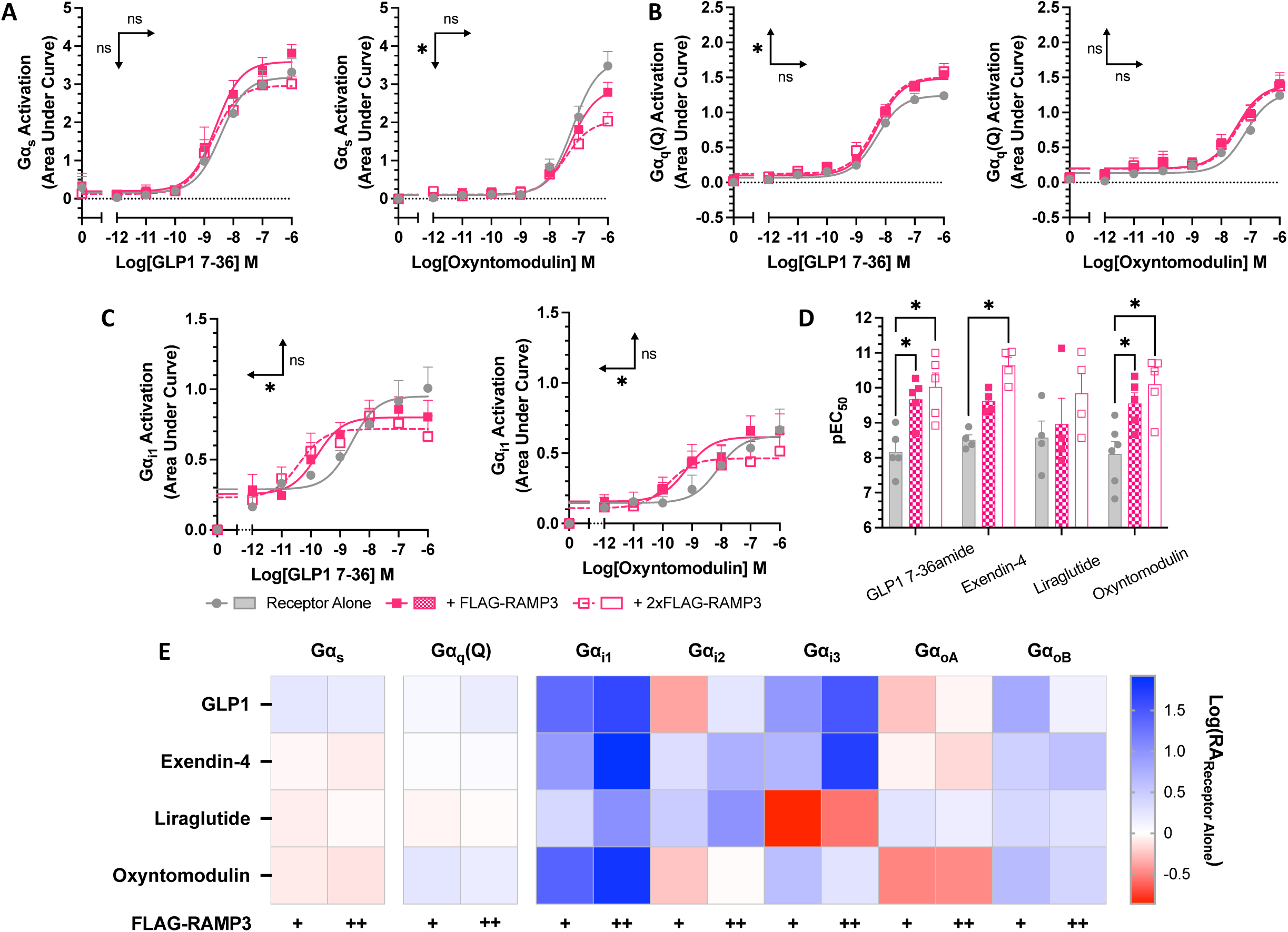
RAMP3 alters G protein coupling of GLP-1R differently depending on the peptide agonist. G protein coupling measured using the TRUPATH biosensor platform in HEK293T cells transfected with GLP-1R. (A) Gα_s_, (B) Gα_q_(R183Q), and (C) Gα_i1_ shown in response to GLP-1 and oxyntomodulin. (D) pEC_50_ values for Gα_i1_ coupling in response to all agonists. (E) LogRA values for all Gα subunits tested, relative to receptor alone. Mean values are shown of n experimental replicates, ± SEM, where n>4. Significance between RAMP conditions determined through One-Way ANOVA, * p < 0.05. Horizontal arrows indicate changes in potency (pEC_50_), and vertical designate changes in efficacy (E_max_).

Whilst the effects on Gα_s_ and Gα_q_provide some insights as to how RAMP3 exerts its effects on second messenger signalling, it is not conclusive. Linked with both a reduction in cAMP, and an increase in (Ca^2+^)_i_ is Gα_i/o_. GLP-1R coupling to this family of G proteins has been observed previously (7, 8), so the effects of RAMP3 were studied across the family. GLP-1R showed little coupling to Gα_z_, which was unchanged by RAMP3 expression. However, for other members of the family, there was a significant increase in the potency of coupling as increasing concentrations of RAMP3 were expressed (Gα_i1_showed in Figure 4C-D, Supporting Table S1).

To better enable comparisons between the different Gα subunits, LogRA values were calculated, relative to the response to receptor alone (Figure 4E, Supporting Table S1). The changes at Gα_s_ and Gα_q_(Q) were slight, mirroring the effects when only looking at the potency (pEC_50_) and efficacy (E_max_) parameters. Larger changes were observed in Gα_i/o_ coupling, with significant decreases in the LogRA for Gα_i1_when RAMP3 was overexpressed in cells stimulated with exendin-4 and oxyntomodulin.

### Signalling in an insulin secreting cell line

GLP-1R signalling is inherently linked with insulin secretion, as receptor activation significantly enhances GSIS. This is linked to both cAMP accumulation and (Ca^2+^)_i_ mobilisation, enhancing sensitivity of the insulin secretion machinery, with β-arrestin recruitment and internalisation both positively and negatively implicated in GSIS (31, 32). To confirm the changes observed in HEK293T cells are applicable to a more physiological setting, the above signalling assays were repeated in INS-1 832/3 cells, a rat insulinoma cell line. These cells endogenously express the GLP-1R, alongside low levels of RAMP2 and 3 (Figure 5A). RAMP3 was transfected into these cells, maintaining the endogenous expression of GLP-1R to look at the second messenger signalling. Assaying at 8- and 30-minute stimulation, no effect was observed on cAMP accumulation in response to GLP-1 when RAMP3 was expressed (Figure 5B, Table 3). However, as observed in HEK293T cells, RAMP3 did increase the potency of (Ca^2+^)_i_ mobilisation (from a pEC_50_ of 6.94±0.11 to 7.41±0.14) (Figure 5C-D, Table 3). When calculating LogRA values, there is bias towards (Ca^2+^)_i_ mobilization, reflecting this enhanced potency (Figure 5E), although the effect was not significant.

**Figure 5.**
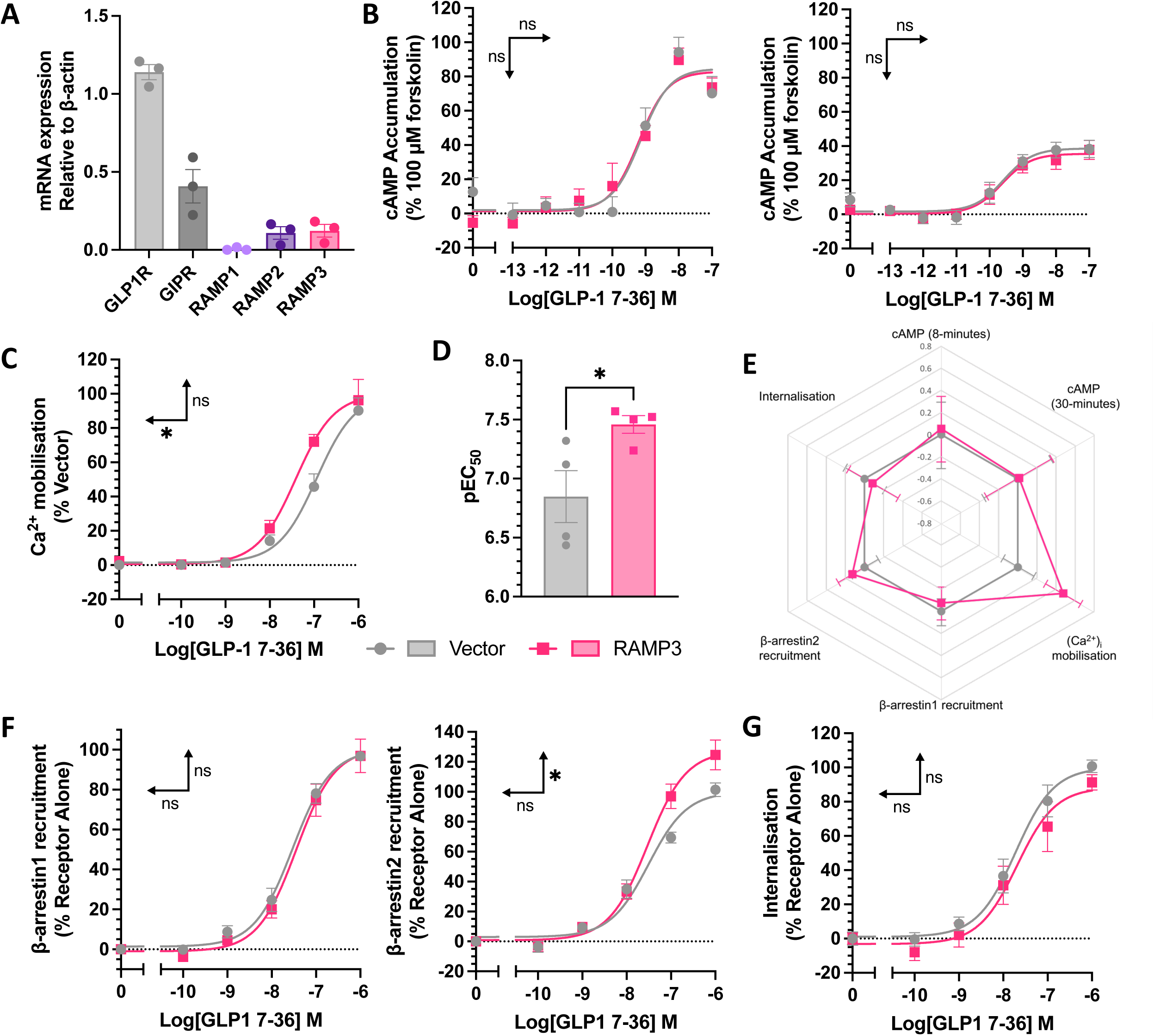
Selective enhancement of Ca^2+^ mobilisation by increased RAMP3 expression. (A) mRNA expression of incretin hormone receptors and RAMPs in INS-1 832/3 cells, relative to β-actin (n=3). (B) cAMP accumulation in response to 8 or 30-minute stimulation with GLP-1, in INS-1 832/3 cells transfected with vector or RAMP3 (n>4). Responses are expressed as percentage of 100μM forskolin (C) Ca^2+^ mobilisation in response to stimulation with GLP-1, in INS-1 832/3 cells transfected with vector or RAMP3, with pEC_50_ values plotted in (D), n=4. Responses are normalised to 10μM ionomycin. (E) Radar plot of LogRA values for second messenger accumulation and receptor desensitisation, relative to receptor alone. (F) β-arrestin1 (*left*) or β- arrestin2 (*right*) recruitment to GLP-1R-Nluc, transiently transfected into INS-1 832/3 cells, n>3. (G) Internalisation of GLP-1R-Nluc in INS-1 832/3 cells, as determined through a loss of BRET with RIT-Venus (n=4). Responses are normalised to vector transfected cells. Mean values of n experimental replicates are shown ± SEM, significance between RAMP conditions determined through Student’s t-test, * p < 0.05. Horizontal arrows indicate changes in potency (pEC_50_), and vertical designate changes in efficacy (E_max_).

When looking at β-arrestin recruitment or internalisation, INS-1 832/3 cells without endogenous GLP-1R expression (33) were used to avoid any potential interference. Cells were transfected with GLP-1R-Nluc and either vector or RAMP3, and β-arrestin1/2-YFP for β-arrestin recruitment, or RIT-Venus for internalisation. RAMP3 had no effect on β-arrestin1 recruitment but increased maximal β-arrestin2 recruitment in this cell line (Figure 5F). This shows a system dependency to the effects of RAMP3, with the opposite effect observed in HEK293T cells. As in HEK293T cells, there was no effect on internalisation (Figure 5G, Table 3).

### Insulin secretion is enhanced by RAMP3

INS-1 832/3 cells overexpressing RAMP3 displayed significantly greater insulin secretion in response to 10nM GLP-1 than those expressing vector; GLP-1 increased insulin secretion 2.36±0.30 fold over 2.8mM glucose compared to 3.94±0.33 when RAMP3 is overexpressed (Figure 6A). This was not observed when cells were transfected with RAMP1, although both RAMP1 and 3 increased insulin secretion in response to 10nM GIP, whose receptor interacts with both proteins (17) (Figure 6A). The effect of RAMP3 was confirmed for the other GLP-1R agonists, whose signalling has also been affected by RAMP3 expression, although the increase was not significant for exendin-4.

**Figure 6.**
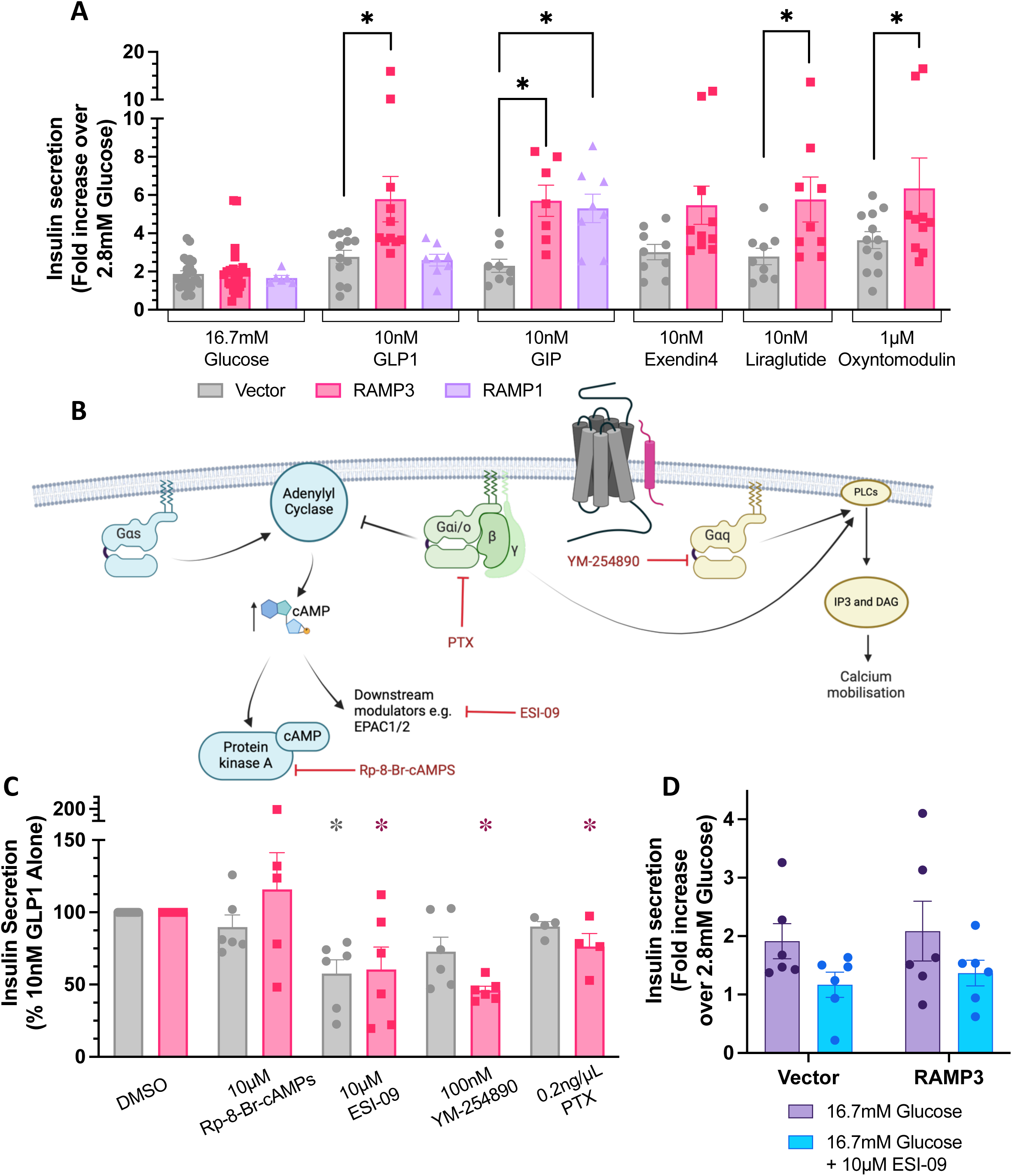
RAMP3 enhances incretin-mediated insulin secretion, dependent on (Ca^2+^)_i_ mobilisation. (A) Glucose-stimulated insulin secretion in INS-832/3 cells, transfected with vector, RAMP3, or RAMP1 and co-stimulated with 16.7mM glucose and GLP-1, GIP, exendin-4, liraglutide, or oxyntomodulin. Results are expressed as fold change over 2.8mM glucose, n>7. (B) Schematic illustrating the action of different small molecule inhibitors of G protein signalling, and their effects on insulin secretion in INS-1 832/3 cells transfected with vector or RAMP3. (C) Insulin secretion in INS-1 832/3 cells transfected with vector or RAMP3 and stimulated with 10nM GLP-1. Cells were pretreated and co-stimulated with inhibitors of G protein signalling. Responses are normalised to cells treated with 10nM GLP-1 and 0.1% DMSO, which acted as a vehicle control, n>4. (D) Insulin secretion in response to 16.7mM glucose, either in the absence or presence of 10μM ESI-09, n=6. Mean values of n experimental replicates are shown ± SEM, significance between RAMP conditions determined through One-Way ANOVA (GIPR) or Two-Way ANOVA (GLP-1R), or Kruskal-Wallis test (GLP-1R inhibitor dataset) * p < 0.05. Horizontal arrows indicate changes in potency (pEC_50_), and vertical designate changes in efficacy (E_max_).

To determine how this increase in insulin secretion occurs, inhibitors of different G protein-mediated signalling pathways were employed (schematic shown in Figure 6B). Data have been normalised to the response to GLP-1 in the absence of inhibitor.

As no specific inhibitor of Gα_s_ is available, inhibitors of proteins whose signalling is downstream of cAMP accumulation were used. Inhibition of Protein Kinase A (PKA) with Rp-8-Br-cAMPs had no effect on insulin secretion, irrespective of the expression of RAMP3 (Figure 6C). However, inhibition of exchange factor directly activated by cAMP1/2 (EPAC1/2) with the dual inhibitor ESI-09 significantly decreased GLP-1 mediated potentiation of GSIS in both vector and RAMP3 transfected cells (Figure 6C). However, a reduction was also observed on the response to 16.7mM glucose alone (Figure 6D).

Inhibition of Gα_q/11_activation with 100nM YM-254890, or Gα_i/o_ signalling with pertussis toxin (PTX) decreased insulin secretion significantly only when RAMP3 was overexpressed. It is therefore suggested that RAMP3 enhances Gα_q/11_and Gα_i/o_ activation, enhancing (Ca^2+^)_i_ mobilisation and GSIS.

### RAMP3 is important for GLP-1R mediated insulin secretion

As these experiments focused on the influence of RAMP3 overexpression on GLP-1R pharmacology, we next investigated the effect of RAMP3 knockdown in circumstances where it is endogenously expressed. Whilst INS-1 832/3 cells express low levels of RAMP3, Min6 B1, a mouse islet cell model was shown to express both RAMP2 and RAMP3. This expression was unaffected by glucose culture conditions, although the GLP-1R expression correlated with increasing glucose concentration (Figure 7A, example gel shown in 7B). RAMP3 mRNA was reduced using siRNA treatment, to around 50% of cells treated with control, scrambled siRNA (Figure 7C). There was no effect on GLP-1R expression, or cAMP accumulation mediated by the GLP-1R or GIPR, following 8-minute stimulation with GLP-1 or GIP respectively (Figure 7D). Insulin secretion mediated by glucose with or without 10nM or 1nM GIP was unaffected by RAMP3 knockdown, but the response to 10nM and 1nM GLP-1 was attenuated, although the effect was small (Figure 1E). This is likely due to the relatively low RAMP3 expression, and only partial knock down in expression.

**Figure 7.**
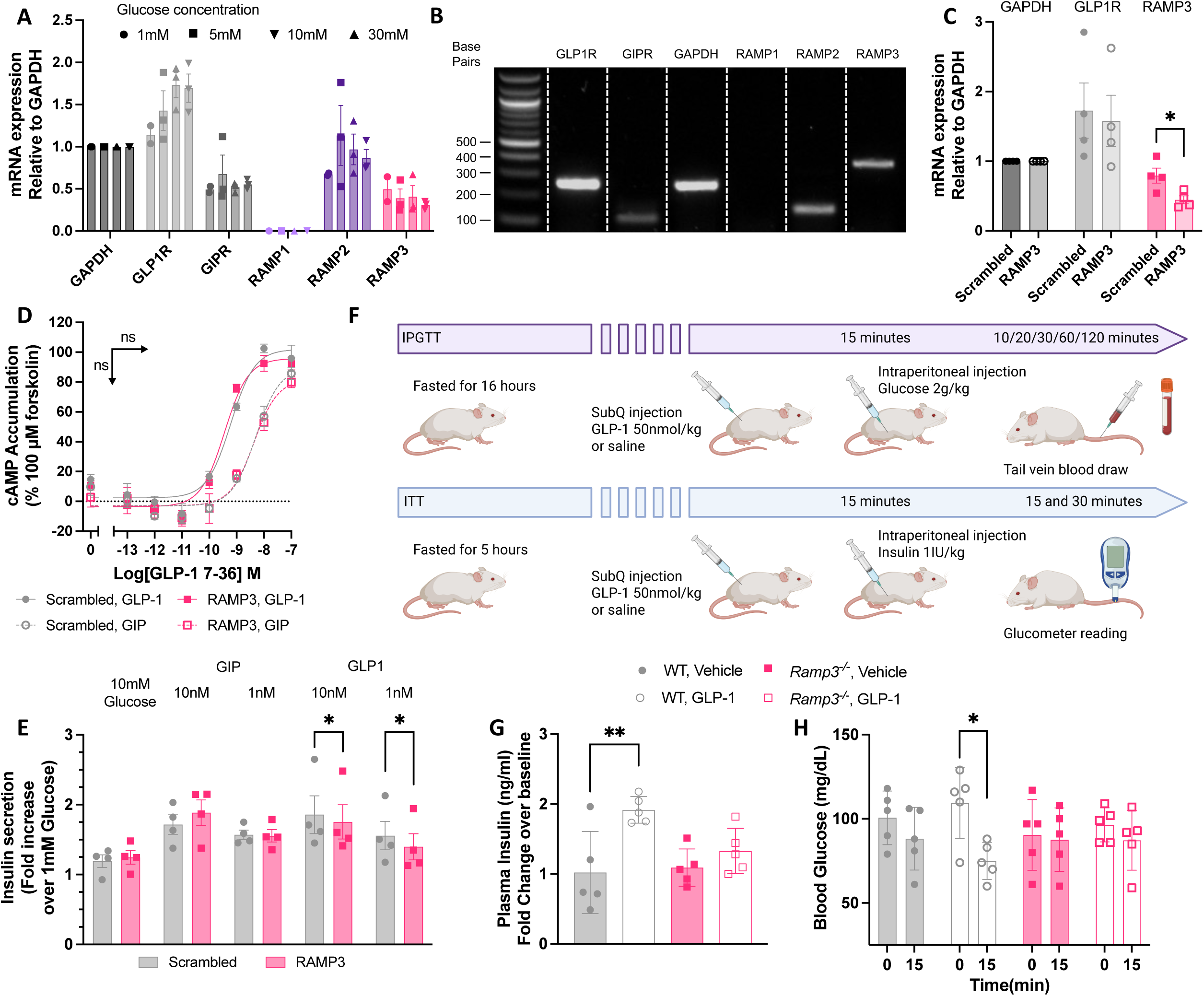
Reducing RAMP3 expression decreases sensitivity to GLP-1 in murine models. (A) mRNA expression of GLP-1R, GIPR, RAMP1, RAMP2, and RAMP3 in Min6 B1 cells, cultured in varied glucose concentrations (n=3), with example gel shown in (B). Data are normalised to GAPDH. (C) mRNA expression of GLP-1R and RAMP3, relative to GAPDH, in Min6 B1 cells treated with scrambled of RAMP3 siRNA (n=4). (D) cAMP accumulation in siRNA treated Min6 B1 cells, in response to 8-minute stimulation with GLP-1 (closed symbols) or GIP (open symbols), n=4. (E) Insulin secretion in response to 10mM glucose, with or without GIP or GLP-1, in Min6 B1 cells treated with scrambled or RAMP3 siRNA, n=4. (F) Schematic for treatment of WT and Ramp3^-/-^ mice for IPGTT and ITT tests. (G) Plasma insulin for IPGTT WT and Ramp3^-/-^ mice, expressed a fold change over baseline responses (n=5). (H) Blood glucose concentrations of WT and Ramp3^-/-^ mice, following ITT. Measured at baseline (0) and after 15 minutes, n=5. Mean values of n experimental replicates are shown ± SEM (or SD for biological repliactes in in G and H). Significance determined through Student’s t-test for RAMP3 knockdown, or matching Two-Way ANOVA (E and H) or One-Way ANOVA (G), with Tukey’s test for multiple comparisons.

To assess how these effects translate into more physiologically relevant systems, we moved to a global Ramp3 gene deletion mouse model (34). This allowed for measurement of intraperitoneal glucose tolerance, (IPGTT) and insulin tolerance (ITT) (Figure 7F). In wild type (WT) mice, GLP-1 increased insulin secretion 15 minutes after administration of glucose. This response was lost in *Ramp3^-/-^*mice. GLP-1 also potentiated the effects of insulin on blood glucose, an effect that too was lost in *Ramp3^-/-^* mice (Figure 7H). Together this data demonstrates an importance of RAMP3 for the functional, rapid response of GLP-1 signalling in the control of glucose homeostasis.

## Discussion

Insulin secretion following ingestion of food high in carbohydrates is greatly potentiated by signalling of incretin hormones. Manipulation of the signalling of the incretins is therefore a valuable strategy for treating diabetes. However, development of therapeutics targeting the GLP-1R has been made difficult by the variability in the signalling of the receptor depending on the study. Part of this variability is attributed to the cell background in which the assay is performed, and the different interacting proteins expressed therein. We therefore sought to identify any interactions with the RAMP family of proteins, which modulate signalling at many Class B1 GPCRs, and determine any subsequent effects on signalling of exogenously and endogenously expressed receptors. We then further delved into the effect of these changes in signalling on GSIS.

GLP-1R was found to interact with RAMP3, but not the other members of the RAMP family of proteins. Previous studies have shown interactions with RAMP1 and 2, although at a reduced level. However, these studies have used isolated lysates (18), or whole cell resonance energy transfer (17, 19). Neither technique discerns between interactions in different subcellular compartments and therefore does not confirm a functional receptor complex. Use of a cell impermeable dye to irreversibly label SNAP-RAMP enables measurement of only interactions at the plasma membrane, suggesting interactions previously observed with RAMP1 and 2 occur in intracellular compartments. Furthermore, adaptation of the NanoBiT system for ligand binding determined it is only the GLP-1R-RAMP3 complex capable of agonist binding, and not the other two RAMPs. This interaction had no effect on membrane expression of either GLP-1R of RAMP3, nor was there any effect on binding of GLP-1 to the receptor. Unlike other RAMPs, RAMP3 is able to translocate to the plasma membrane in the absence of an interacting GPCR (12). It is therefore possible that it displays interactions with an increased number of GPCRs through proximity, as opposed to a specific interaction (26). However, increasing the expression of RAMP1 at the membrane through overexpression of CLR had no effect on the interaction observed.

RAMP3 expression did, however, modulate G protein signalling at the receptor. Consistent with previous observations (19), cAMP accumulation was decreased in the presence of RAMP3, especially at early time points. However, unlike previous reports, an increase in (Ca^2+^)_i_ mobilisation was observed with increasing expression of RAMP3. These findings were validated by examining the effects on G protein coupling, using the TRUPATH biosensor platform. At individual agonists, small perturbations could be observed. Gα_s_ coupling in response to oxyntomodulin was increased by RAMP3, and Gα_q_coupling was increased when cells were stimulated with GLP-1. However, for the remaining agonists, no effects were observed. This did not correlate with the second messenger signalling results, where effects were more widespread across the agonists. Gα_i/o_coupling, whilst small, was increased by RAMP3 in response to all agonists tested. In correlation with the effects on cAMP production, β-arrestin recruitment was decreased by RAMP3 overexpression, although not for the weaker agonist oxyntomodulin. There were no subsequent effects on internalisation for most agonists, but the response to liraglutide was reduced. The GLP-1R has been shown to internalise through multiple avenues, with no β-arrestin dependency (27), so a lack of correlation is not surprising.

In many of the assays studied, ligand-dependent effects were observed e.g. selective enhancement of different G proteins, or differential effects on efficacy compared to potency. The agonists themselves have their own inherent signalling bias, with liraglutide displaying bias towards β-arrestin recruitment. This may then correlate with arrestin-dependent internalisation not observed for the other agonists, explaining why RAMP3 only alters internalisation in this case. It is thought that agonist bias stems from stabilisation of distinct receptor conformations, promoting different G protein couplings (35). RAMPs can also alter GPCR conformational states (36), leading to different effects depending on the ligand. Without structural interrogation, it is difficult to ascertain how these agonist-dependent effects emerged. However, these differences were primarily the magnitude of the effect, with RAMP3 still generally moving GLP-1R towards (Ca^2+^)_i_ mobilisation and insulin secretion in response to all agonists.

Insulin secretion is enhanced by overexpression of RAMP3. This was observed for all GLP-1R agonists tested, but not high glucose alone. It is therefore unlikely the increase in RAMP3 expression itself directly increases insulin secretion. Furthermore, increasing RAMP1 expression (which does not interact with GLP-1R) had no effect on the potentiation of GSIS by GLP-1R, but did enhance GIPR mediated GSIS, consistent with findings that reducing RAMP1 expression reduced glucose tolerance in mice (17).

GLP-1-mediated GSIS was unaffected by inhibition of PKA by Rp-8-Br-cAMPs in the presence or absence of RAMP3. Previous studies have shown potentiation of GSIS to be both PKA dependent (37) and independent (38), with this study agreeing with the latter, although this might be due to reduced affinity of the compound for the rat kinases. Inhibition of EPAC1/2 with ESI-09 significantly reduced insulin secretion, although a non-significant reduction was also observed in cells stimulated with high glucose alone, suggesting importance in GSIS independent of the incretin hormones. EPAC1/2 has been previously shown to be important for cAMP-dependent potentiation of GSIS (39), which this data supports. As cAMP accumulation was unaffected in the INS-1 832/3 cell line by the overexpression of RAMP3, it is not surprising that there was no interactive effect of RAMP3 when PKA or EPAC1/2 were inhibited. YM-254890 and PTX, inhibitors of Gα_q/11_ and Gα_i/o_ respectively, however, only reduced insulin secretion in cells overexpressing RAMP3. This is consistent with the elevation in (Ca^2+^)_i_ mobilisation when RAMP3 was overexpressed, dependent on enhanced Gα_i/o_ coupling. Gα_i/o_-mediated (Ca^2+^)_i_ mobilisation has been shown to depend on active Gα_q/11_ (40), making the resolution of the respective contributions of the two G protein families difficult. However, the effect of YM-254890 was greater than PTX, suggesting a contribution from both families.

In murine models, endogenous RAMP3 was shown to be important, as a reduction in expression attenuated effects of GLP-1 on insulin secretion and glucose tolerance. As with the overexpression data for INS-1 832/3 cells, RAMP3 knockdown had no effect on cAMP accumulation. It is possible this difference is due to lower receptor expression, or a differential effect on bias for the rodent receptors than the human. Species differences in signalling bias have been observed for other GPCRs associated with islet cell pharmacology (41), so it stands to reason this might be influencing the differences observed here. However, with over 90% sequence identity, it is more likely that the effects seen in the HEK293T cells is a product of the receptor overexpression, as this has been noted as a factor in agonist bias (42). When expanded to an animal model with global deletion of *Ramp3*, this reduction in insulin secretion became more pronounced, with GLP-1 no longer able to elevate insulin secretion above vehicle-treated animals. This then led to an elevation in blood glucose concentration, again through a loss of response to GLP-1. *Ramp3* knockout has previously been shown to increase the effect of GLP-1 on bodyweight lowering in mouse models, highlighting the preservation of a functional receptor within these animals (43) This also highlights the complexity of GLP-1R pharmacology, with tissue and context dependent effects of *Ramp3* KO; an elevation in one function, but a loss of another. These differences may stem from the deletion of only *Ramp3* in this study compared to the dual *Ramp1/Ramp3* deletion described in Leuthardt *et al*., or may relate to differences in GLP-1R/RAMP3 pharmacology in endocrine vs neuronal tissues. These results may therefore present an opportunity to further separate the role of the receptor in glucose homeostasis and appetite regulation.

Within this study, RAMP3 has been shown to influence agonist bias at the GLP-1R, shifting the receptor towards (Ca^2+^)_i_ mobilisation, which enhances insulin secretion. As RAMP3 expression varies between tissues, it is possible that by directly targeting this complex, tissue specific receptor activation could be facilitated. However, of more relevance is the consideration of RAMP expression when developing new GLP-1 mimetics, or small molecule agonists and allosteric modulators for the treatment of T2DM and obesity. Agonists which have reduced affinity for the RAMP3 complex over receptor alone might show reduced therapeutic potential, which might not be detected when only looking at the GLP-1R alone. Furthermore, it is important to ensure allosteric modulators do not disrupt the interaction, as is the case for the CLR-RAMP1 antagonist BIBN 4096 BS (44), as this could lead to a reduction in GLP-1R activity in cells expressing RAMP3.

## Materials and Methods

### Materials

GLP-1 (7-36amide), oxyntomodulin, exendin-4 and GLP-1 (9-36amide) were custom synthesised by Generon. Liraglutide was purchased from Cayman Chemicals, and GIP and GLP-1 (for *in vivo* experiments) were purchased from Bachem. Peptides were dissolved at 1mM (or 10mM in the case of GLP-1 9-36amide) in DMSO, or water with 0.1% BSA *w/v* for GIP and GLP-1 (9-36amide). Forskolin was purchased from Cayman Chemicals and made up to 10mM in DMSO, and ionomycin was purchased from Cayman Chemicals and made up to 10mM in ethanol. YM-254890 (Wako Chemicals, Alpha Laboratories) was made up to 1mM, and Rp-8-Br-cAMPs (SigmaAldrich) and ES109 (Bio-Techne) were stored at 10mM in DMSO. PTX (Gibco, Thermo Scientific) and stored as a 0.1mg/mL solution in water (*w/v*).

### DNA Constructs

LgBiT-GLP-1R and Nluc-GLP-1R were made by PCR amplification of GLP-1R and restriction enzyme digest (Invitrogen, Thermo Scientific). This was then ligated into pcDNA3.1(+)sig-LgBiT or pcDNA3.1(+)sig-Nluc (where sig is the signal peptide of the murine serotonin 5-HT3R). GLP-1R-Nluc was made via the same process but was ligated into pcDNA3.1(-)-Nluc. SmBiT-RAMPs were generated by restriction enzyme digest of the RAMP with EcoRI and EcoRV (Invitrogen, Thermo Scientific), the product of which was then ligated into pcDNA3.1(+)-sigSmBiT. Sequences were confirmed by Sanger sequencing (University of Cambridge). RIT-Venus was donated by Luke Pattison (Jimenez-Vargas et al., 2018), and β-arrestin1/2-YFP were donated by Duncan Mackie (Mackie et al., 2019). Untagged RAMP1 and RAMP3 were purchased from cDNA.org. Use of Nluc-CLR, SNAP-RAMPs, and FLAG-RAMP3 has been described previously (17, 45). TRUPATH G protein constructs were a gift from Bryan Roth (Addgene kit #1000000163).

### Cell Culture and transfection

Cos7 and HEK293T cells were grown in Dulbecco’s Modified Eagle Media (DMEM)/F12 Glutamax supplemented with 1% Antibiotic-Antimycotic (AA) solution and 10% Foetal Bovine Serum (FBS) *v/v*. INS-1 832/3 and INS-1 832/3 GLP-1RKO cells, donated by Jacqueline Naylor (AstraZeneca), were grown in RPMI supplemented with 1% AA, 5% FBS, 10mM HEPES, 50μM β-mercaptophenol, and 1mM sodium pyruvate. Min6 B1 cells (gifted by Dr. Philippe Miyazaki, University of Osaka) were cultured in DMEM containing 4.5g/L glucose, supplemented with 15% FBS, 1% penicillin-streptomycin (PS), and 50μM β-mercaptophenol. Cells were grown at 37°C with 5% CO_2_. Transfection of cells with polyethyleneimine (PEI) was conducted at a 6:1 *v/w* ratio of PEI:DNA in 150mM NaCl. Transfection with FuGENE HD (Promega) was carried out according to the manufacturer’s guidance, using a 3:1 *v/w* ratio of FuGENE HD:DNA. Transfection of INS-1 cells lines was carried out with Lipofectamine 2000 according to the manufacturer’s instructions, using a 3:1 ratio *v/w* of Lipofectamine 2000:DNA. For siRNA knockdown of RAMP3, Min6 B1 cells were treated with 20nM siRNA for 48 hours, using lipofectamine 3000 at a 3:1 *v/w* ratio. Control scrambled siRNA was purchased from Invitrogen (Negative control siRNA 1). ON-TARGETplus SMARTpool siRNA targeting mouse RAMP3 was purchased from Horizon Discovery (Revvity), containing sequences: 5’-GGUGCAACCUGUCGGAGUU-3’, 5’-GUAUGCGGCUGCAACGAGA-3’, 5’-GCACCGAGAUGGAGACCAA-3’, 5’_-_GAAGUACUCAUCCCACUGA-3’.

### Receptor-RAMP interactions and binding

Experiments were adapted from those detailed in Harris et al., 2021. Cos7 cells were seeded at 15,000 cells per well of a 0.01%-poly-L-lysine coated 96-well white Culturplate and incubated overnight. Cells were transfected with 50ng Nluc-GPCR and increasing amounts of SNAP-RAMP. After 24 hours, growth media was replaced with serum free DMEM/F12 containing 200nmol of SNAP-Surface Alexa Fluor-488 and incubated for 30 minutes. Cells were washed and incubated in KREBs (126 mM NaCl, 2.5 mM KCl, 25mM NaHCO3, 1.2 mM NaH2PO4, 1.2 mM MgCl2, 2.5 mM CaCl2) containing 0.1% Bovine Serum Albumin (BSA) *w/v* and 0.1% Nano-Glo® Cell Assay Substrate (Promega) *v/v* and read using a Mithras LB microplate reader at 460nm and 515nm. The ratio at the 10-minute time point was used for further analysis. For NanoBiT interaction assays, HEK293T cells expressing increasing amounts of SmBiT-RAMP, and 100ng LgBiT-GPCR were incubated in KREBs containing 0.2% NanoGlo® Cell Assay Substrate *v/v*. The 10-minute time point was used to construct saturation curves.

The Taglite GLP-1R fluorescent probe (Cisbio), based on exendin-4, was used for binding assays (Ex-4-Red). LgBiT-GLP-1R and SmBiT-RAMP, or Nluc-GLP-1R and FLAG-RAMP3 were co-expressed in HEK293T cells at a 1:1 or 1:2 ratio. Cells were incubated in KREBs containing 0.1% or 0.2% *v/v* NanoGlo for 5 minutes. Immediately before reading, Ex-4-Red in the presence of GLP-1, 10μM exendin-9, or vehicle was added. Where injection was used, 50μl of 1μM GLP-1 in KREBs supplemented with 0.1% BSA (*w/*v) and 0.1% Nano-Glo® (*v/*v) was injected robotically at 20-minutes into a well containing 50μl KREBs supplemented with 0.1% BSA (*w/*v) and 0.1% Nano-Glo® (*v/*v) and 4nM Ex4-Red. Plates were read using a Mithras LB 940 multimode microplate reader, using 460nm and 610nm long pass emission filters at 30 second intervals.

### Flowcytometry to measure GPCR and RAMP Surface Expression

For GLP-1R surface expression, HEK293T cells were transfected with Nluc-GLP-1R and FLAG-RAMP3 at a 1:1 or 1:2 ratio. After 48 h, 300,000 cells were washed twice with FACS buffer (PBS supplemented with 1% BSA and 0.03% sodium azide) before and after incubation with rabbit anti-Nluc polyclonal antibody, diluted 1/100 (Promega) for 1 h at room temperature in the dark. Samples were analysed using a BD Accuri C6 flow cytometer, Ex. λ 633 nm and Em. λ 660 nm. Data were normalised to the median APC intensity of cells transfected with pcDNA3.1 as 0% and Nluc-GLP-1R + pcDNA3.1 as 100%. For RAMP cell surface expression, HEK293T cells were transfected with Nluc-GLP-1R or HA-CLR and FLAG-RAMP1/2/3 at a 1:1 ratio. After 48 h, 300,000 cells were washed twice with FACS buffer (PBS supplemented with 1% BSA and 0.03% sodium azide) before and after incubation with rat allophycocyanin (APC)-conjugated anti-Flag monoclonal antibody for 1 h at room temperature in the dark. Data were normalized to the median APC intensity of cells transfected with pcDNA3.1 as 0% and HA-CLR + FLAG-RAMP2 as 100%.

### Reverse transcriptase PCR to measure endogenous GPCR and RAMP expression

For studies assessing the effects of glucose culture conditions, Min6 B1 cells were cultured in glucose-free DMEM, supplemented with 15% FBS, 1% PS, 50μM β-mercaptophenol and 1mM, 5mM, 10mM, or 30mM glucose for 4 days prior to harvest. RNA was extracted from INS-1 832/3 and Min6 B1 cells using RNeasy mini kit (Qiagen) as per the manufacturer’s protocol. Complementary DNA was generated using the QuantiTect reverse transcription kit (Qiagen) following the manufacturer’s instructions with minus Reverse Transcriptase negative controls performed simultaneously. PCR amplification was performed as previously described (20, 46) using the following gene-specific primers: rat β-actin, forward (5′-CCGCGAGTACAACCTTCTTG-3′) and reverse (5′-CAGTTGGTGACAATGCCGTG-3′); rat GLP-1R, forward (5′-GGGCTCCTCTCGTATCAGGA-3′) and reverse (5′-GTGAACAGCTTGACGAAGCG-3′); rat GIPR, forward (5′-AGGTGGTATTTGCTCCCGTG-3′) and reverse (5′-AGGGGTCCCTTTACCTAGCA-3′); rat RAMP1, forward (5′-GATGTGAGGACAGGAACCAGA-3′) and reverse (5′-TGGTCTTTCCCCAGTCACAC-3′); rat RAMP2, forward (5′-CTCCGGAGTCCCTGAATCAA-3′) and reverse (5′-TCCAGTTGCACCAGTCCTTG-3′); rat RAMP3 forward (5′-ACAAACATCGTGGGCTGCTA-3′) and reverse (5′-CCACGGTCAACAAGACTGGA-3′); mouse GAPDH forward (5’-CAGGAGAGTGTTTCCTCGTCC-3’) and reverse (5’-GATGGGCTTCCCGTTGATGA-3’); mouse GLP-1R forward (5’-ACTCTCATCCCCCTTCTGGG-3’) and reverse (5’-GGACACTTGAGGGGCTTCAT-3’); mouse GIPR forward (5’-CGAGTGGCCAGAGTTTCCAT-3’) and reverse (5’-TCTGCCCCTCAGAGTCTGTC-3’); mouse RAMP1 forward (5’-AGCCGCTTCAAGGAGAACAT-3’) and reverse (5’-CGAAACTGCTTCCTGCAAACT-3’); mouse RAMP2 forward (5’-TGAGGACAGCCTTGTGTCAA-3’) and reverse (5’-GGTCGCTGTAATGCCTGCTA-3’); mouse RAMP3 forward (5’-AGTACTTCAGAGCCTAGAGGTGA-3’) and reverse (5’-ATAGCCACAGTCAGCACGAC-3’). Products were resolved on a 2% agarose gel and imaged using an iBright imager (Invitrogen). Band intensity was analysed using Fiji (ImageJ) and normalised to the β-actin (INS-1 832/3) or GAPDH (Min6 B1).

### cAMP Accumulation

HEK293T cells and INS-1 832/2 cells were cultured as before and transfected 24 hours before assaying, with Nluc-GLP-1R, and pcDNA3.1 and/or FLAG-RAMP3 (HEK293T) or 48 hours before with pcDNA3.1 or RAMP3 (INS-1). Min6 B1 cells were seeded for 48 hours following siRNA treatment. On the day of assay, cells were dissociated into single cell suspension with Trypsin 0.05% EDTA and resuspended in phosphate buffered solution (PBS) with 0.1% BSA *w/v* in the presence (INS-1 832/3 and Min6 B1) or absence (HEK293T) of 500μM IBMX. Cells were plated at 1000 (HEK293T), 2000 (INS-1 832/3), or 500 (Min6 B1) cells per well of a 384-well Optiplate (PerkinElmer) in 5μl aliquots. Cells were stimulated for a given period and then cAMP accumulation measured using the LANCE *ultra* cAMP detection kit as per the manufacturers’ instructions. Data were normalised to 100 μM forskolin.

### Calcium Mobilisation Assay

HEK293T cells were transfected with 100ng Nluc-GLP-1R and 200ng of pcDNA3.1 or FLAG-RAMP3, or 100ng of pcDNA3.1 and FLAG-RAMP3 per 100μl cells and seeded in a 0.01% PLL-coated black-walled clear-bottom 96-well plate. 48 hours later media was removed, and cells were incubated in 10μM Fluor4-AM dye containing 2.5mM probenecid for 1 hour at room temperature. Cells were washed and dye replaced with Hank’s Balanced Salt Solution (Lonza) without Ca^2+^ containing 0.1% BSA *w/v*. Ligands were added robotically using a BD Pathway 855 high intensity bioimager, and images taken at 0.5 second intervals for 120 seconds. The response to vehicle was taken to be 0%, and the response to 10 μM ionomycin was used as 100%. For HEK293T cells, the maximal fluorescence was used for further analysis. For INS-1 832/3 cells, the area under the curve for the first 120 seconds after ligand addition was used.

### G Protein Dissociation

G Protein dissociation assays were performed using the TRUPATH biosensor platform, as described (28). HEK293T cells were transfected with GLP-1R, Gα, Gγ, Gβ, and pcDNA3.1 or FLAG-RAMP3 all in equal quantities and incubated overnight. Cells were then seeded on white 96-well 0.01% PLL-coated plates in MEM media supplemented with 1% AA and 2% FBS at 50,000 cells per well and cultured overnight. Cells were washed with KREBs with 0.1% BSA and incubated in buffer containing 5μM coelenterazine 400a for 5 minutes. Ligands were then added, and plates read using the Pherastar microplate reader using the BRET2 module, at 60-second intervals. The BRET2 ratio was calculated (λ_515_/λ_400_) and total dissociation over a 30-minute period used to generate dose response curves. Data were normalised to the response of receptor alone to GLP-1 at each Gα protein.

### β-arrestin recruitment assays

β-arrestin recruitment assays were adapted from those previously described (22, 47). Cells were transfected with GLP-1R-Nluc, β-arrestin-YFP and either pcDNA3.1 or FLAG-RAMP3 in a 1:5:2 ratio and incubated overnight. HEK293T cells were then reseeded onto a 0.01% PLL-coated white 96-well Culturplate at 50,000 cells per well. INS-1 832/3 cells were reseeded at 100,000 cells per well in growth media. Stimulation buffer for HEK293T cells was KREBs containing 0.1% BSA, whereas KREBs-Ringer (2.6 mM CaCl_2_, 98.5 mM NaCl, 4 mM KCl, 1.2 mM KH_2_PO_4_, 1.2 mM MgSO_4_, 20 mM HEPES, 25.9 mM NaHCO_3_, pH 7.4) with 0.1% BSA was used for INS-1 832/3 cells. Cells were washed before being incubated in stimulation buffer with 0.1% Nano-Glo® Cell Assay substrate for 5 minutes. Ligand was then added, and plates were immediately read using a Mithras LB 940 microplate reader, at 460nm and 530nm, for 1 hour at 60 second intervals. Data were normalised the response of receptor alone to GLP-1 at each arrestin.

### Measurement of Receptor Internalisation

Experiments were performed as previously described ((47)). Briefly, HEK293T or INS-1 832/3 GLP-1R KO cells were transfected with GLP-1R-Nluc, RIT-Venus and either pcDNA3.1 or FLAG-RAMP3 in a 1:5:2 ratio and incubated overnight. HEK293T cells were then reseeded onto a 0.01% PLL-coated white 96-well Culturplate at 50,000 cells per well, in MEM media supplemented with 1% AA and 2% FBS. INS-1 832/3 cells were reseeded at 100,000 cells per well in growth media. Stimulation buffer for HEK293T cells was KREBs containing 0.1% BSA, whereas KREBs-Ringer (2.6 mM CaCl_2_, 98.5 mM NaCl, 4 mM KCl, 1.2 mM KH_2_PO_4_, 1.2 mM MgSO_4_, 20 mM HEPES, 25.9 mM NaHCO_3_, pH 7.4) with 0.1% BSA was used for INS-1 832/3 cells. Cells were washed with stimulation buffer before being incubated in stimulation buffer with 0.1% Nano-Glo® Cell Assay substrate for 5 minutes. Ligand was then added, and plates were immediately read using a Mithras LB 940 microplate reader, at 460nm and 530nm, for 1 hour at 60 second intervals. Data were normalised the response of GLP-1R alone to GLP-1.

### Insulin Secretion

INS-1 832/3 cells were transfected with pcDNA3.1 or RAMP3 and seeded at 75,000 cells per well of a 0.01% PLL-coated clear 96-well plate. Min6 B1 cells were seeded at 50,000 cells per well following siRNA treatment. For experiments using PTX, after 24 hours PTX was added to a final concentration of 0.2ng/μL and maintained throughout the assay. 48 hours after transfection, the cells were glucose starved in RPMI media without glucose (INS-1 832/3) or DMEM media with 1g/L glucose (Min6 B1) for 3 hours. Cells were washed and incubated at 37°C for 1 hour in stimulation buffer (KREBs-Ringer with 0.1% BSA) containing 2.8mM (INS-1 832/3) or 3mM (Min6 B1) glucose, before being stimulated for 1 hour. Where signalling inhibitors were used, cells were incubated with the inhibitor or vehicle control for 30 minutes prior to stimulation. The supernatant was removed, and the cells incubated for 5 minutes in stimulation buffer with 1% Triton-X 100 (v/v). Supernatant and lysed cell solution were centrifuged at 2000rpm for 5 minutes and diluted. Insulin content was measured with the insulin ultra-sensitive detection kit (Cisbio). Values are expressed as fold change over low glucose.

### Mouse intraperitoneal glucose tolerance test (IPGTT) and insulin tolerance test (ITT)

All animal studies were approved by the Institutional Animal Care and Use Committee of UNC-Chapel Hill. Animal housing, care, and husbandry were overseen by the UNC Division of Comparative Medicine Animal Resources, which is accredited by the International Association for Assessment and Accreditation of Laboratory Animal Care (AAALAC). Prior to experiments, mice were multi-housed in a temperature-controlled room with a 12:12 dark light cycle and provided ad libitum access to water and standard chow.

To measure the glucose lowering effect of GLP-1(7–36) peptide mice IPGTT and ITT were performed on wildtype SvEv129/SJ and isogenic *Ramp3^-/-^* male animals. *Ramp3^-/-^*mice and genotyping have been previously described (34). For IPGTT, mice were fasted for 16 hr and treated with saline or GLP-1 at 50nmol/kg body weight (subQ) 15 min before glucose (2g/kg body weight) intraperitoneal injection. Blood glucose was measured using a glucometer (True Matrix, Trividia Health, Inc) from the tail vein at baseline (before GLP-1 injection),10, 20, 30, 60 and 120 min after glucose injection. Blood was collected at baseline and 15 min after glucose injection from the tail vein in EDTA cuvettes (Microvette CB 300, 16.444.100, Sarstedt). For ITT, mice were fasted for 5 hr and treated with saline or GLP-1 at 50nmol/kg body weight (subQ) 15 min before insulin (1IU/kg) intraperitoneal injection. The blood glucose levels were measured from the tail vein at baseline (before GLP-1 injection),15 and 30 min after insulin injection using a glucometer.

### Detection of in vivo plasma insulin levels

During IPGTT, blood samples were collected in potassium EDTA cuvettes from mouse tail vein at baseline and 15 min after glucose administration. Blood samples were centrifuged at 2000g for 10 min at 4°C to separate the plasma (supernatant) which was collected in separate 1.5 ml Eppendorf tubes and were stored at −80°C. Using the mouse insulin ELISA kit (Ultrasensitive mouse insulin ELISA kit, Crytal Chem, 90080), plasma insulin levels were measured following the manufacturer’s instructions with two replicates for each sample.

### Data Analysis

GraphPad Prism 10 was used for analysis of all assays. cAMP, Ca^2+^ mobilisation, β-arrestin recruitment, and internalisation were fitted to a concentration-response curve using the three-parameter logistic equation. Potency (pEC_50_) and maximal (E_max_) values were obtained from this curve fit. Saturation-interaction curve data was fitted using a one-phase decay model and binding data fitted using either a model of association kinetics, or the equation for a one site binding model. LogRA values were calculated using

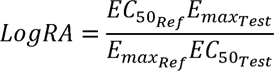

Where Ref is the reference condition, and Test that for which the LogRA is being calculated.

Error was propagated as previously described ((46, 48).

Statistical differences were determined using a Student’s t-test, One-Way or Two-Way ANOVA, with Dunnet’s, Tukey’s, or Sidak’s post-hoc test for multiple comparisons for parametric data, or Kruskal-Wallis test with Dunn’s post-hoc test for multiple comparisons for non-parametric datasets. A probability of p<0.05 was considered significant. Values are reported as mean ± SEM of n repeats carried out in duplicate or quadruplicate.

## Acknowledgments

A.P. is funded by the Biotechnology and Biological Sciences Research Council (BBSRC; BB/JO14540/1; BB/W014831/1). H.Y.Y. was supported by a Cambridge Trust International Scholarship and the Rosetrees Trust. C.M.S was funded by a Cambridge Trust International Scholarship in conjunction with the Gonville and Caius Stanley Elmore PhD Studentship. S.W. was supported by an AstraZeneca studentship. G.L. is a Royal Society Industry Fellow (INF/R2/212001). P.K. and K.M.C are supported by NIH/NICHD HD060860, and acknowledge the technical assistance of Ms. Megan Brantly.

## Data Availability

All data is contained within the manuscript.

